# Reproducible Bioconductor Workflows Using Browser-based Interactive Notebooks and Containers

**DOI:** 10.1101/144816

**Authors:** Reem Almugbel, Ling-Hong Hung, Jiaming Hu, Abeer Almutairy, Nicole Ortogero, Yashaswi Tamta, Ka Yee Yeung

**Affiliations:** Institute of Technology, University of Washington, Tacoma, WA, USA; Department of Clinical Investigation, Madigan Army Medical Center, Tacoma, WA, USA

**Author notes:** Co-first authors (Reem Almugbel, Ling-Hong Hung). Corresponding author: Ka Yee Yeung.

**Keywords:** Bioconductor workflows, containers, reproducibility, automated, data science

## Abstract

**Objective:** Bioinformatics publications typically include complex software workflows that are difficult to describe in a manuscript. We describe and demonstrate the use of interactive software notebooks to document and distribute bioinformatics research. We provide a user-friendly tool, BiocImageBuilder, to allow users to easily distribute their bioinformatics protocols through interactive notebooks uploaded to either a GitHub repository or a private server.

**Materials and methods:** We present three different interactive Jupyter notebooks using R and Bioconductor workflows to infer differential gene expression, analyze cross-platform datasets and process RNA-seq data. These interactive notebooks are available on GitHub. The analytical results can be viewed in a browser. Most importantly, the software contents can be executed and modified. This is accomplished using Binder, which runs the notebook inside software containers, thus avoiding the need for installation of any software and ensuring reproducibility. All the notebooks were produced using custom files generated by BiocImageBuilder.

**Results:** BiocImageBuilder facilitates the publication of workflows with a point-and-click user interface. We demonstrate that interactive notebooks can be used to disseminate a wide range of bioinformatics analyses. The use of software containers to mirror the original software environment ensures reproducibility of results. Parameters and code can be dynamically modified, allowing for robust verification of published results and encouraging rapid adoption of new methods.

**Conclusion:** Given the increasing complexity of bioinformatics workflows, we anticipate that these interactive software notebooks will become as ubiquitous and necessary for documenting software methods as traditional laboratory notebooks have been for documenting bench protocols.

## BACKGROUND AND SIGNIFICANCE

Bioinformatics is an interdisciplinary research area focused on developing and applying computational methods derived from mathematics, computer science, and statistics to analyze biological data [1]. Workflows typically involve the execution of a series of computational tasks. Documenting workflows has become increasingly difficult given the growing complexity of workflows and rapidly evolving methodologies. Traditional “Materials and Methods” sections are not well suited for describing methodologies that are strongly dependent on code and parameters. Recently, public software repositories such as GitHub have made it relatively straightforward to distribute and update code. Data science notebooks such as Jupyter are the software analogs to laboratory notebooks and document the parameters and code along with the results. Formal descriptors of workflows such as Common Workflow Language [1] that describe how different software components interact are also gaining acceptance. While these steps go a long way to documenting computational workflows, it is estimated that more than 25% of computational workflows cannot be reproduced [2].

The problem is that even when using the correct version of each code component, executing them in the correct order, with the correct parameters on identical data, is still not sufficient to ensure that the same results are obtained [3]. Running the same software in a different hardware and software environment can affect the outcome. Even if this were not the case, from a practical viewpoint, these dependencies can make installation of the exact version of the same components problematic on a different system. Using software suites such as Bioconductor [4] has become a popular method to manage multiple packages and ensure proper installation and interoperability. However, the rapid development of new tools has made it increasingly difficult to define a base setup that is compatible with the growing number of components that are potentially written in different programming languages.

A solution to this problem is to package software inside software computers that provide all the requisite software dependencies. Docker is a very popular software container technology and has been used for large scale deployment and re-analyses of biomedical big data [5]. JupyterHub [6] allows users to run Jupyter notebooks inside software containers deployed locally or through a server. Binder [7] takes this a step further and generates and deploys Docker images of notebooks from GitHub on their public servers. This allows users to view and interact with notebooks in GitHub repositories *without* the need to compile code or install software. However, using Binder or Jupyter requires writing custom Dockerfiles to specify the elements present in the Docker container.

In this paper, we demonstrate the utility of live notebooks for documenting and distributing bioinformatics workflows by presenting three notebooks on GitHub that use Bioconductor. In addition, we present a framework and a graphical user interface (GUI) designed to automatically generate a Dockerfile for a custom Bioconductor installation. This allows a user without any technical knowledge of Docker, to generate and publish live Bioconductor notebooks.

### Reproducibility of Bioinformatics workflows using Bioconductor

Reproducibility is essential for verification and advancement of scientific research. This is true for computational analyses, which are not reproducible in more than 25% of publications [2]. Reproducibility in bioinformatics research refers to the ability to re-compute the data analytics results given a dataset and knowledge of the data analysis workflow [8]. For this to happen, three requirements must be available: (i) datasets (ii) code and scripts used to perform the computational analyses (iii) metadata; details about how to obtain and process datasets, including description of software and hardware environment setup [9 10]. Gentleman and Lang proposed a compendium software framework for the distribution of dynamic documents containing text, code, data, and any auxiliary content to re-create the computations [11]. Their framework forms the foundation of the Bioconductor project [4] that provides an online repository for software packages, data, metadata, workflows, papers and training materials, as well as a large and well-established user community. Bioconductor also works with the broader R repository, the Comprehensive R Archive Network (CRAN) [12] which also contains useful bioinformatics packages that are not included with Bioconductor.

### Using Software Containers to Enhance Reproducibility of Bioinformatics Workflows

Unfortunately, re-running published code and data to reproduce published results is non-trivial even when using Bioconductor. Bioconductor is not static: new packages are constantly being added and other packages deprecated. Correct versioning is thus essential for reproducibility. Although Bioconductor is cross-platform, it achieves this by cross-compilation which does not completely insulate the computation from differences in local hardware and software. The solution is to run the software in a virtual environment that is the same regardless of the underlying hardware or host operating system.

Docker is a well-established container technology to increase reproducibility of bioinformatics workflows [3 10 13]. The Docker platform allows for virtualized application deployments within a lightweight, Linux-based wrapper or container [14]. Essentially, containers are virtual environments that encapsulate only the minimum necessary dependencies, which can be quickly deployed on most major platforms in a reproducible fashion [14]. Docker uses a Dockerfile that contains all the instructions to build a Docker image starting from scratch or from another Docker image. Docker images can be downloaded from repositories like Docker Hub (<u>https://hub.docker.com/</u>).

While Docker provides an easy, modular method to build, distribute and replicate complex pipelines and workflows across multiple platforms, widespread adoption in the biomedical field has been hampered by the level of technical knowledge presently required to use the technology. Docker was developed for computer professionals with programming and systems administration experience who are able to write a Dockerfile to script the installation of the software environment for a container. Our group has worked on enabling graphical interfaces to interact with containers to make Docker more accessible to less technical users [10 15 16].

### Data Science Notebooks

All laboratories use notebooks to document the experimental procedures and protocols. The software counterparts are data science notebooks that combine text, code, data, mathematical equations, visualizations, and rich media into a single document that can be accessed through a web browser. These software notebooks first gained popularity in mathematics research, and the Jupyter open source project has expanded the scope/audience to include many heavily computational research areas such as bioinformatics, neuroscience and genomics [17]. Project Jupyter was a spinoff project from IPython that supports the Python programming language and now maintains multiple kernels for over 50 programming languages including Ruby, Javascript, C++ and Perl [18 19]. Each Jupyter notebook document is divided into individual cells that can be run independently [18]. This format records every step of a computational analysis along with the scientific narrative, which makes it easier to understand, share, reproduce and extend a published computational workflow.

To facilitate notebook sharing and reusability, Jupyter project supports *nbconvert*, a tool to convert notebooks to various formats such as HTML, PDF and LaTex [20]. It also supports *nbviewer*, a similar web service to view and download any Jupyter notebook publically published on the web [21]. In 2015, GitHub (<u>https://github.com/</u>), a web-based version control code repository, started supporting Jupyter notebook format by making it possible to render Jupyter notebooks written in any programming language on GitHub [22], which brings GitHub’s features of sharing, version control and collaboration into the Jupyter platform. Today, there are more than 940,000 Jupyter notebooks rendered on GitHub [23].

### Sharing live notebooks using Binder

While *<u>static</u>* Jupyter notebooks can be shared and viewed using a browser *<u>without any setup</u>* via *nbviewer* and GitHub, sharing *<u>interactive dynamic</u>* Jupyter notebooks in which the user can execute and modify the analyses requires the notebooks to be downloaded and installation of Jupyter. To address this limitation, the Binder open source project (<u>http://mybinder.org/</u>) offers a browser-based executable environment to run Jupyter notebooks hosted on GitHub. The Binder environment allows scientists to share *live interactive* Jupyter notebooks that are reproducible and verifiable using a web browser, with no data download or software installation requirements [17]. To manage computational environments, Binder’s underlying architecture takes advantage of two open source projects: Docker, which builds the environments from a project’s dependencies, and Kubernetes, which schedules resources for these environments on a Google Compute Engine cluster [24]. To build and launch an executable binder, a Jupyter notebook must be uploaded to a public GitHub repository, along with an environment specification such as a Dockerfile [7]. Scientists have used Binder as a publishing medium to share reproducible computational workflows [25]. However, most of the use cases of Binder have been limited to the iPython kernel. An example is the Laser Interferometer Gravitational-Wave Observatory (LIGO) that used iPython Jupyter notebooks and Binder to demonstrate the computational workflow corresponding to the first direct detection of gravitational waves that Einstein predicted decades ago [26 27].

Although many bioinformatics workflows use R and Bioconductor, the use of interactive notebooks has been mostly restricted to Python-based workflows due to the difficulties in setup. The software dependencies required for Binder must be reconciled with the strict dependencies required for the base installation R and Bioconductor. Any customization steps must also be included in the setup files. To enable more widespread adoption of interactive notebooks in bioinformatics, we need a more accessible method to prepare an interactive notebook as provided by BiocImageBuilder.

## OUR CONTRIBUTIONS

We present a novel software tool, BiocImageBuilder, that *automates* the technical step of creating Dockerfiles for live Jupyter notebooks using Bioconductor. A web-based graphical interface allows the user to choose the Bioconductor packages that need to be installed and whether the Dockerfile is to be used with Binder on GitHub or is to be used in a local installation. Additional R packages from CRAN can also be included. The tool then builds the appropriate Dockerfile for the user to upload with their notebook. We illustrate the feasibility of our tool in three case studies: differential expression analyses for ectopic pregnancy, pattern discovery of gene expression data across human cell lines and a published RNA sequencing workflow. All of our case studies use the R programming language and software packages from Bioconductor/CRAN. Figure 1 shows an overview of our approach.

**Figure 1.**
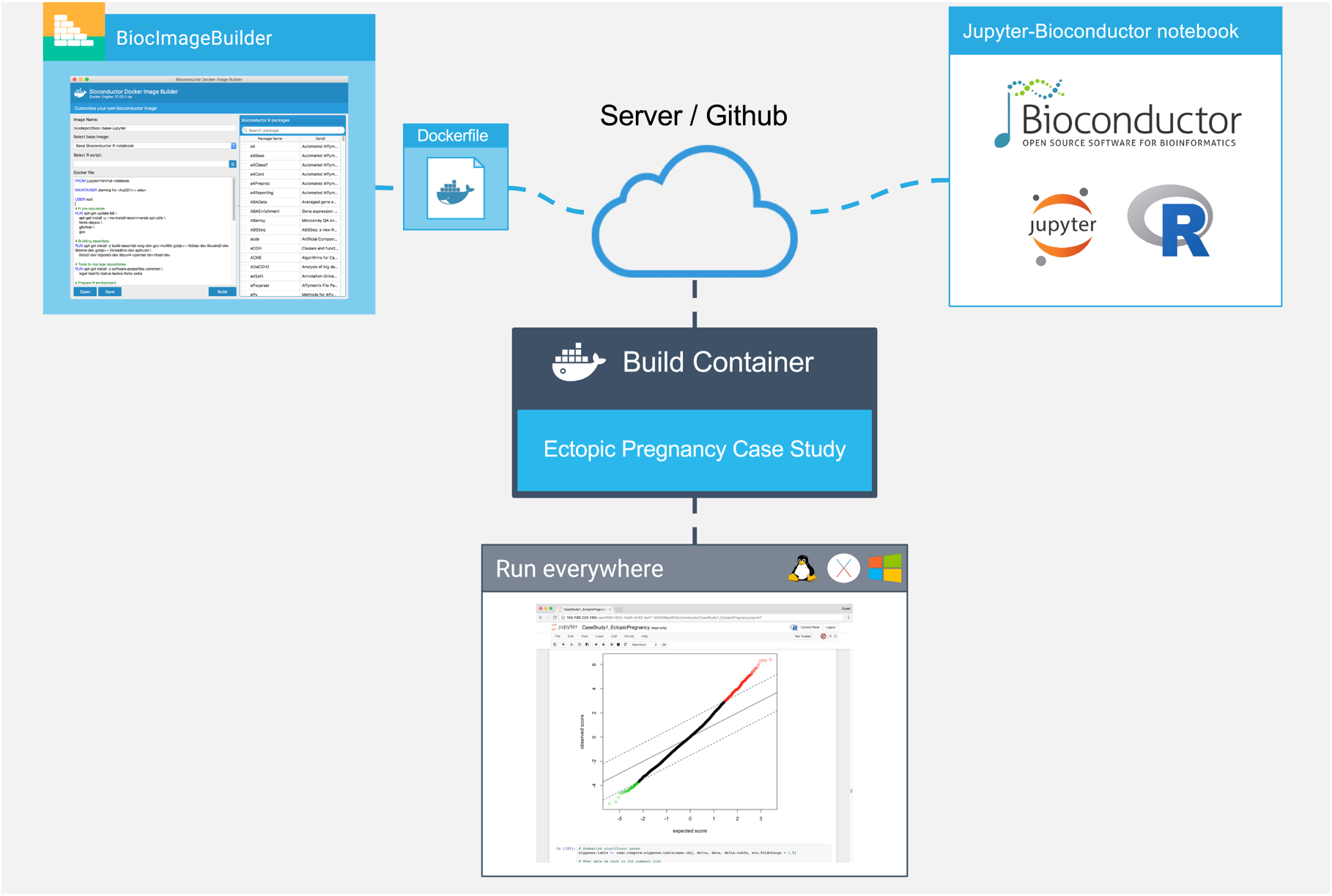
Overview of our approach. The author of the Bioconductor workflow uses BiocImagebuilder to generate a Dockerfile that describes the Bioconductor and CRAN packages installed. The Dockerfile and the notebook files are uploaded to a server or GitHub repository. A custom container is then built with the default Linux base-image for Bioconductor, dependencies for Jupyter, JuptyerHub and/or Binder and the Bioconductor packages. For GitHub installations, the Binder server builds the container and provides a link to run the container on their public cluster. JupyterHub provides the same functionality locally or on a private server. Using the container, the end-user is able to view the notebook, execute, modify and save the code on their local machine regardless of whether it uses Linux, MacOS or Windows. In the case, where the container is run remotely, no additional installation of software is required on the part of the end-user.

## AUTOMATIC GENERATION OF DOCKERFILES FOR BIOCONDUCTOR WORKFLOWS

Bioconductor compiles and tests each component of its suite on a set of stock Windows, MacOS and Linux machines. This ensures that all components in Bioconductor are compatible and should install on most hardware configurations. In addition, the Bioconductor core team provides Docker containers for the release and development versions of the complete suite [28]. No facility exists however, for building custom images with a specified set of Bioconductor components.

We have developed a GUI-based tool, BiocImageBuilder, for this purpose. BiocImageBuilder starts with a base-image that is based on the stock Linux test machine. The base-image is modified to include components for Binder compatibility if desired, and the kernels necessary to run R. For images to be run by Binder, the Linux Conda utility is used to install the Bioconductor and CRAN packages. Otherwise, Bioconductor’s biocLite utility is used install the components. The user simply starts up the container with BiocImageBuilder and points their browser to a local URL. They will then see a form for choosing the desired starting image, the desired components and the option of running a custom startup script (see **Figure 2** and **Supplementary File 1**). BiocImageBuilder will then produce a Dockerfile. This can be uploaded to GitHub, along with a Jupyter notebook file to create a repository that distributes an interactive notebook that can be viewed using Binder. Alternatively, a Dockerfile can be produced that is suitable for private deployment using JupyterHub. Users can also use the Dockerfile to directly build an actual image themselves of their notebook to use, store or distribute on DockerHub and other repositories. Currently, we support R 3.4 and Bioconductor 3.5. We intend to add support for other versions in the future so that deprecated packages can be run. Containerizing Jupyter notebooks ensures that they will always be viewable, insulating the user from future changes to Bioconductor or R.

**Figure 2.**
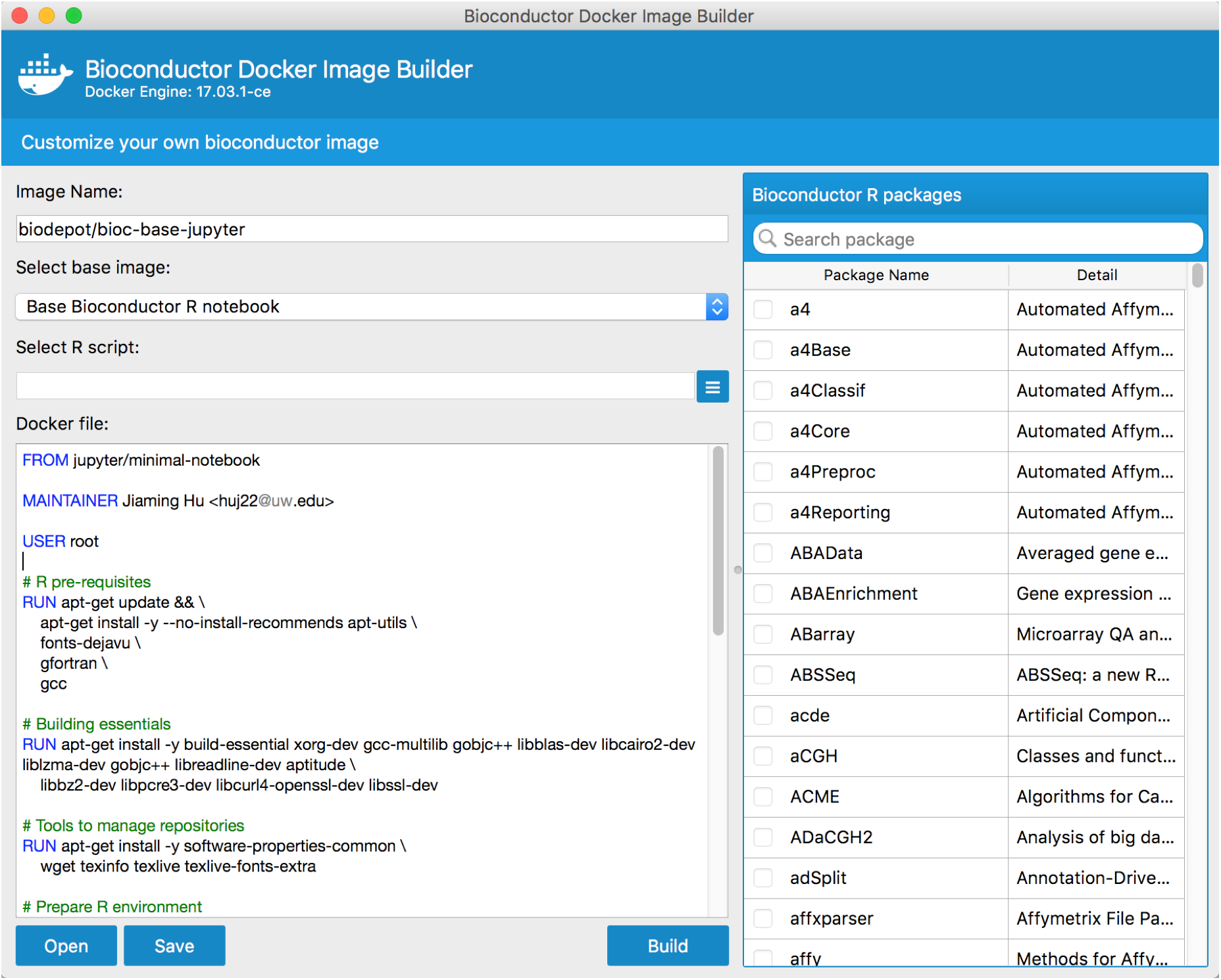
Screenshot of BiocImageBuilder. The user selects from a menu the Bioconductor and Cran packages required for their notebook. BiocImageBuilder then generates the Dockerfile describing a minimal Linux container that contains these packages. The Dockerfile can be uploaded to GitHub where it can be viewed interactively using Binder.

BiocImageBuilder is written in Python3 using PyQt5 (<u>https://wiki.python.org/moin/PyQt</u>) which is a Python binding for the Quicktime engine that renders the graphical interface. Although PyQt5 is meant to be cross-compatible over different platforms, there are many dependencies and installation can be quite complicated for some user environments. To avoid these problems, BiocImageBuilder is packaged using our GUIdock-noVNC container [15]. This container creates a mini-webserver that serves the rendered graphics through a local port, and can be run on any Docker compatible platform (Windows, MacOS, Linux). Most modern browsers that support HTML5 (e.g. Chrome, Firefox, Safari, Opera) can be used to access the BiocImageBuilder. Note that BiocImageBuilder is designed for those wishing to author an interactive Bioconductor notebook - it is not required for end users wishing to interact with a published notebook. The source code of BiocImageBuilder is publicly available at <u>https://github.com/Bioconductor-notebooks/BiocImageBuilder</u> and its Docker image is publicly available at <u>https://hub.docker.com/r/biodepot/bioc-builder/</u>.

## CASE STUDIES

In this section, we present three case studies in which we illustrate the use of R and Bioconductor packages within Jupyter notebooks. Static snapshots of these notebooks are included as Supplementary Files 2-4. The corresponding fully interactive notebooks are available from <u>https://github.com/Bioconductor-notebooks</u>. BiocImageBuilder was used to automatically generate the necessary Dockerfiles for Binder. In Case Study 1, we extended the published differential expression analyses of ectopic pregnancy. In Case Study 2, we created our own workflows for cross-platform omics data. In Case Study 3, we replicated a published RNA-seq workflow in our proposed framework.

### Case Study 1: Identification of differentially expressed genes for ectopic pregnancy

#### Motivation and overview

When a woman’s pregnancy test result is positive, initial testing of the uterus is visualized on a transvaginal ultrasound scan (TVS). As shown in **Figure 3**, the possible outcomes of the TVS are: (i) Intrauterine pregnancy (IUP) which is the case of normal pregnancy with fertilized egg implanted inside the uterus (ii) Ectopic Pregnancy (EP) where the fertilized egg can be seen in the TVS scan, but it is implanted outside the uterus (iii) Pregnancy of Unknown Location (PUL) when the pregnancy test is positive but no evidence of pregnancy is seen on TVS [29].

**Figure 3.**
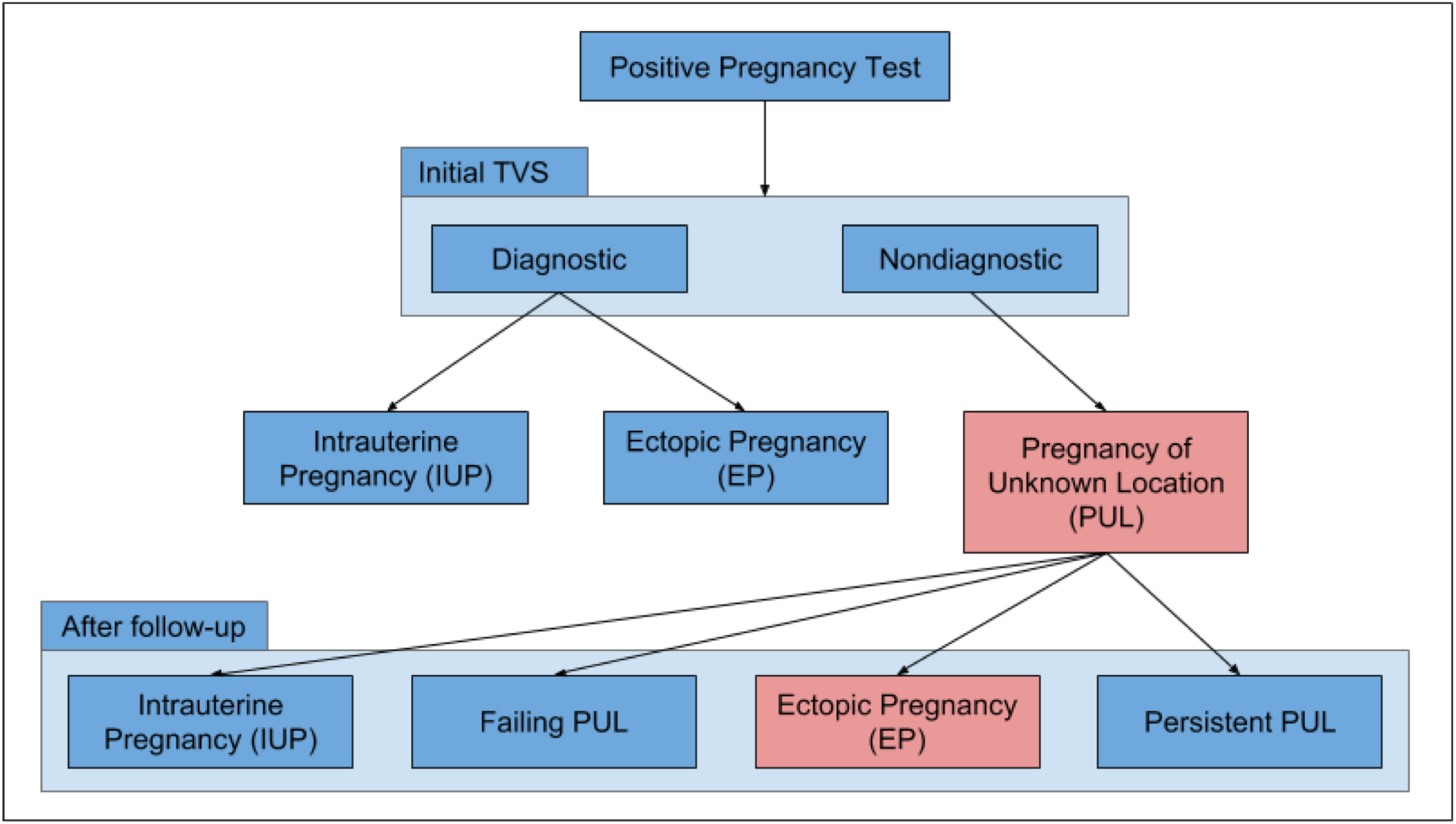
Outcome of initial TVS scan. PUL = Pregnancy of unknown location; TVS = Transvaginal ultrasound scan; EP= Ectopic pregnancy.

Cases of pregnancy of unknown locations (PUL) can subsequently lead to one of the following outcomes: (i) Failing PUL (miscarriage): majority of cases (50-70%) (ii) Normal IUP: fertilized egg is too early to be visualized on TVS (iii) Ectopic pregnancy: 7-20% of the PUL cases, the EP was not seen on the initial TVS examination [29 30].

In the case of PUL, close surveillance is required, consisting of serial office visits, ultrasounds and blood draws over a period as long as a six weeks [31]. During this surveillance period, no medical or surgical intervention is taken until a conclusive diagnosis of ectopic pregnancy is reached, and the non-viability of the embryo is concluded [31]. Thus, the clinicians’ objectives are to: (1) Diagnose ectopic pregnancy as early as possible to avoid health risks, (2) Ensure that this early diagnosis is correct, to avoid ending a viable pregnancy erroneously [32]. Delayed diagnosis of EP is the most common life-threatening emergency in early pregnancy [31]. Despite the high frequency of this serious condition, early diagnosis of EP can be challenging. In practice, there are several methods used to detect EP in the case of PUL, and they largely depend on biochemical markers such as serum progesterone levels [33] and serum human chorionic gonadotrophin (hCG) levels [34]. However, the biochemical markers used are not consistent [35], and the International Society of Ultrasound in Obstetrics and Gynecology is encouraging the use of mathematical models to expedite EP detection [36]. In this case study, we aim to identify differentially expressed genes among patients with EP by analyzing of gene expression data. Differentially expressed genes are the subset of genes that exhibit expression patterns associated with a EP medical condition.

#### Data

Duncan *et. al* collected gestation-matched endometrium from women with EP (n = 11) and intrauterine pregnancies (IUP) (n = 13), and samples were profiled using the Affymetrix Human Genome U133 Plus 2.0 platform [37]. The CEL files were normalized using RMA (Robust Multiarray Average) [37], and are publicly available from ArrayExpress (<u>http://www.ebi.ac.uk/arrayexpress</u>) with accession number E-MTAB-680.

#### Analysis

We filtered the RMA normalized gene expression data to keep the probe sets that are common with prospective validation samples profiled using Affymetrix genechip Human Gene 2.0 ST. AnnotationDbi [38] and Stringr [39] Bioconductor packages were used to access, map, and process gene identifiers in specific chip annotation databases [40 41]. Duncan *et al*. identified genes differentially expressed in EP versus IUP using the t-test with multiple comparison correction using the Benjamini-Hochberg false discovery detection method with a corrected P-value of <0.05 [37]. In our analysis, we started by performing a standard t-test without corrections, with a range of varying threshold values. We also performed other multiple test correction methods including the Bonferroni correction, SAM [42], and LIMMA [43]. Our resulting lists of differentially expressed genes showed considerable overlap with the results from Duncan *et al*. In particular, Duncan’s top up-regulated gene CSH1 resulted from most of our differential expression analyses, and Duncan’s top down-regulated gene CRISP3 resulted from SAM analysis and Benjamini-Hochberg detection method. We also generated heatmaps to visualize these differentially expressed genes. We observed that EP and IUP samples were mostly assigned to distinct clusters with the exception of two IUP samples that clustered with EP samples, which Duncan *et al*. referred it to the effect of decidualization degree. Furthermore, we performed Gene Set Enrichment Analysis (GSEA) [44] to identify pathways and functional categories among the differentially expressed genes. The details of the analyses are provided in Supplemental File 2.

### Case Study 2: Cross-platform analyses of human cell line genomics data

#### Motivation and overview

The LINCS (Library of Integrated Network based Cellular Signatures) program, funded by the National Institutes of Health, generates different types of data, including gene expression, proteomic, and cell imaging data, in response to drug and genetic perturbations (<u>http://lincsproject.org/</u>) [45]. One of the main objectives of the LINCS program is to study gene signatures resulted from perturbations applied to human cell lines. In particular, the LINCS L1000 gene expression data measure the expression level of approximately 1000 landmark genes in response to drug and genetic perturbation experiments across multiple human cell lines. We aim to study the similarity patterns in the L1000 data across different cell lines. The LINCS L1000 gene expression data are publicly available from the Gene Expression Omnibus (GEO) database with accession number <u>GSE70138</u>. Our goal is to study the consistency of cell line similarities across the LINCS L1000 data and other data sources. In particular, we used the LINCS L1000 gene expression data to explore similarities between different cell lines using different analysis methods, including clustering and dimension reduction techniques. Our work is inspired by Zhang *et al* [46] in which multiple datasets, including the Cancer Cell Line Encyclopedia (CCLE) data [47] and Cancer Genome Project (CGP) data [48] were used to explore the similarity of cell lines and drugs. The results of this study suggested that similar cell lines are expected to have similar drug responses, and similar drugs are expected to have similar effects on a cell line.

#### Data

We used the L1000 data processed by the L1K++ pipeline, an alternative data processing pipeline for the L1000 gene expression data, that we developed at the University of Washington Tacoma. L1K++ is implemented in C++ using linear algorithms to make it over 1000x faster than the available pipelines [49]. We substantiated our results from L1K++ processed data using published cell line gene expression data generated using microarray and RNA-sequencing technology. The Cancer Cell Line Encyclopedia (CCLE) gene expression data used Affymetrix microarrays to profile the genome-wide transcription activities across approximately 1000 human cancer cell lines [47]. The CCLE data is publicly available from the GEO database with accession number <u>**GSE36133**</u>. Similarly, Klijn *et al*. [50] used RNA-sequencing technology to profile the expression across 675 untreated human cancer cell lines. This data is publicly available from ArrayExpress database with accession number E-MTAB-2706 <u>https://www.ebi.ac.uk/arrayexpress/experiments/E-MTAB-2706/</u>.

#### Analysis

In the Jupyter notebook (see Supplementary File 3), we read the three datasets (L1K++, CCLE, RNAseq) including all cell lines and genes. Then, we standardized each one of the three datasets separately by computing the z-score for gene expressions across all cell lines. In order to compare the results from the L1K++ data to those from the other two datasets, we first computed the intersection of genes and cell lines in common between L1K++ and CCLE, which resulted in 55 cell lines and the landmark genes. For L1K++ and RNAseq, we found 41 cell lines in common. Subsequently, we calculated the pairwise distances (including Euclidean distances and squared Mahalanobis distances) and correlation coefficients (including Pearson’s correlation and rank-based Kendall’s correlation) between each pair of cell lines based on their gene expression profiles. We then applied hierarchical clustering and model-based clustering [51] to cluster L1K++ vs. CCLE and L1K++ vs. RNA-seq cell lines.

### Case Study 3: Alignment and differential analyses of RNA-seq analysis workflows

#### Motivation and overview

With the rapidly decreasing costs of sequencing technology, RNA sequencing (RNA-seq) has become a well-established technology to measure gene expression. Here, we demonstrate the feasibility and merits of using an interactive Jupyter notebook to document a published RNA-seq data analyses workflow in Bioconductor [52] (<u>https://www.bioconductor.org/help/workflows/rnaseqGene/</u>).

#### Data

We used the RNA-seq data from the Bioconductor “airway” package in which airway smooth muscle cells were treated with dexamethasone, a synthetic glucocorticoid steroid with anti-inflammatory effects [53]. Glucocorticoids are used, for example, by patients with asthma to reduce inflammation of the airways. In the experiment, four primary human airway smooth muscle cell lines were treated with 1 μM dexamethasone for 18 hours. For each of the four cell lines, we have a treated and an untreated sample. The data are also publicly available in the GEO database with accession number GSE52778.

#### Analysis

We followed the steps of analyses published by Love *et al*. [52]. In particular, we started with the BAM files that provide the alignment data in a binary format. After normalizing the table of read counts, we performed differential expression analyses using DESeq [54], visualization using heatmaps for sample distances, and a mean average plot for the estimated model coefficients.

## DISCUSSION

We present a web-based framework and a graphical user interface (GUI) designed to automatically generate a Dockerfile to create and publish live R and Bioconductor notebooks for bioinformatics workflows without any technical knowledge of Docker containers. These web-based notebooks can be published and viewed with modifiable and executable code in a browser without the installation of any software. We demonstrate the applications of these interactive notebooks using three case studies in which we show the revolutionary aspects of dynamic live notebooks compared to traditional static reports and visualization for data analysis. Notebooks generated in our framework ensure reproducibility of analyses through the use of software containers. Our interactive notebooks enable clinicians and biomedical scientists to visually interact with the analyses while exploring the results through different types of interactive visualizations (e.g. Plotly [55] in case study 2). In addition, parameters can be modified easily. Our approach and BiocImageBuilder is <u>not</u> limited to bioinformatics applications that use Bioconductor software packages, but can be used for any applications that use the R programming language and software packages from CRAN.

A limitation of Jupyter notebooks is that each notebook is limited to one kernel supporting a single programming language. All of our three case studies used the IRkernel that assumes a R programming environment. However, modern bioinformatics workflows consist of modules that are potentially written in different programming languages. For future work, we would like to extend these notebooks to allow for a modular structure consisting of different computing environments.

## SUPPLEMENTARY FILES

**Supplementary File 1:** BiocImageBuilder demo video, publicly available at <u>https://youtu.be/HftUChnYytw</u>

**Supplementary File 2:** Full executable version of the notebook is available at: <u>https://github.com/Bioconductor-notebooks/Identification-of-Differentially-Expressed-Genes-for-Ectopic-Pregnancy</u>

**Supplementary File 3:** Full executable version of the notebook is available at: <u>https://github.com/Bioconductor-notebooks/Cross-platform-Analyses-of-Human-Cell-Line-Genomics-Data</u>

**Supplementary File 4:** Full executable version of the notebook is available at: <u>https://github.com/Bioconductor-notebooks/Dynamic-Re-analysis-RNA-seq-differential-expression-workflow</u>

## FUNDING

L.H.H., and K.Y.Y. are supported by National Institutes of Health grant U54-HL127624. Almugbel and Almutairy gratefully acknowledge the full sponsorship from the Saudi Arabian Cultural Mission (SACM) scholarship program 2015–2017.

## AUTHOR CONTRIBUTIONS

R.A. drafted the manuscript and was primarily responsible for figuring out how Binder works. J.H. implemented and wrote documentation for BiocImageBuilder. L.H.H. added the noVNC container to BiocImageBuilder and refined the container. K.Y.Y. designed and coordinated the study. R.A., A.A. and Y.T. created the Jupyter notebooks for case study 1, 2 and 3 respectively. R.A. and N.O. performed the analyses of the ectopic pregnancy data. J.H., L.H.H. and R.A. created the figures. J.H. created the movie uploaded as Supplementary File 1. All authors tested BiocImageBuilder and edited the manuscript.

## ACKNOWLEDGEMENT

We would like to thank Mr. Yuvi Panda and Mr. Chris Holdgraf from the Berkeley Institute of Data Science for providing access to the latest version of Binder.

